## Supplementary Material for "Capturing cell heterogeneity in representations of cell populations for image-based profiling using contrastive learning"

#### A. Additional results for testing the generalizability of CytoSummaryNet

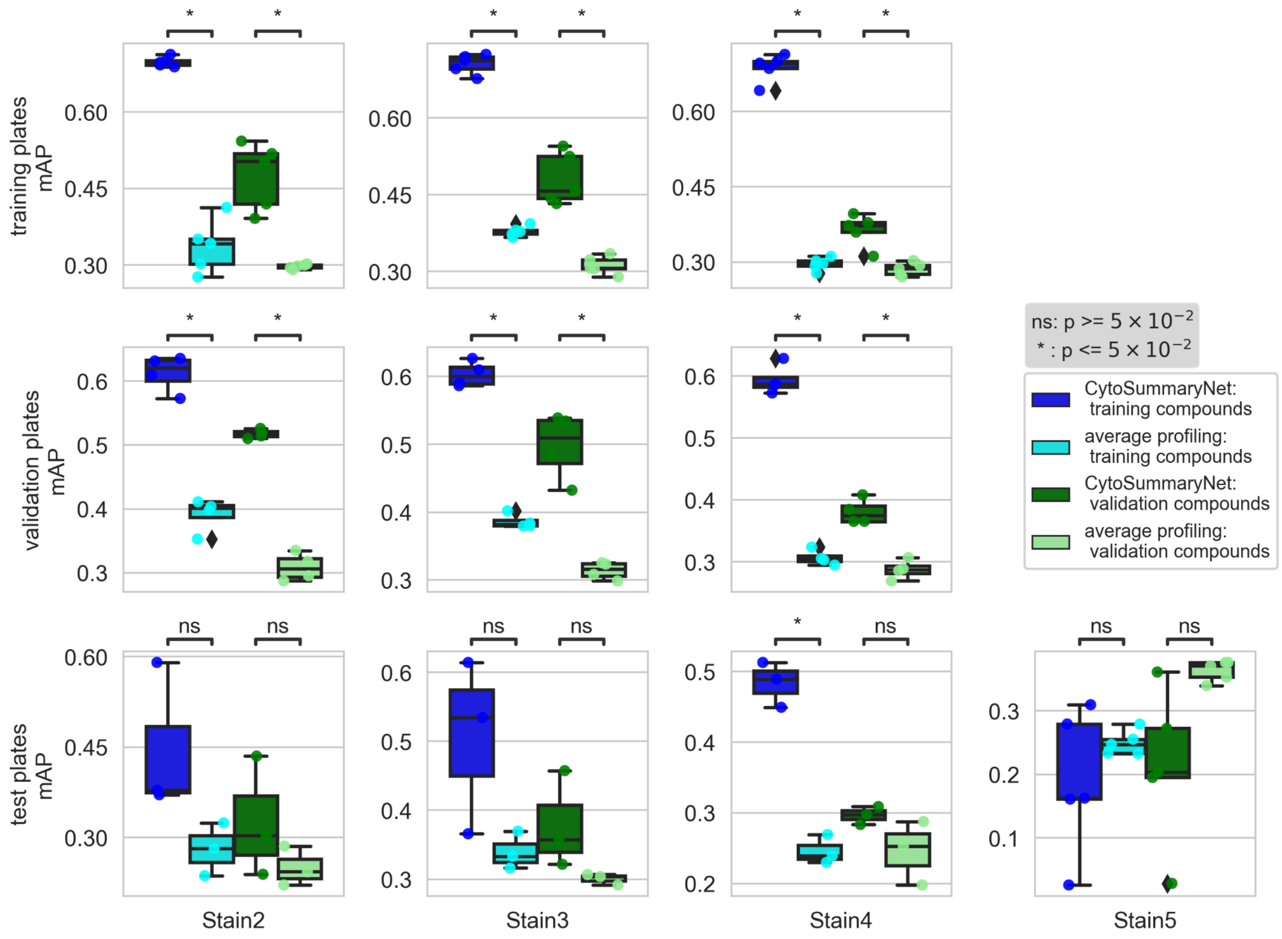


*Figure A1: CytoSummaryNet (model) profiles partially generalize to unseen compounds and do not generalize to unseen experimental conditions. The figure is an expanded version of Figure 4, showing the training and validation plates in addition to the test plates. mAP boxplots of replicate prediction for training and validation compounds by the model- (dark blue and green respectively), and average- (light blue and green respectively) aggregated profiles. mAP scores were calculated separately per training, validation, and test set (rows) and per experiment dataset Stain2, Stain3, Stain4, and Stain5 (columns). Each data point is the average mAP of a plate. Welch's t-tests were used to compare the means between the model and average mAP scores on corresponding data; their p-values are indicated as stars at the top of each plot.*

#### B. Estimating second- and third-order moments using Deep Sets and contrastive learning

A series of experiments was conducted to test if the Deep Sets model, as proposed in the study, can identify different populations based on their second- or third-order moments. In the first experiment, a toy dataset of $n$ different populations was used. The mean of each population was subtracted and the model was trained to distinguish the different populations. The second experiment repeated this setup, but used the real single-cell feature data instead, as described in the Experimental Setup. A third experiment was then conducted with the toy dataset to test the degree to which the model can learn to infer second-order moments from the input data. This was achieved by first subtracting the mean of the populations and then gradually sphering them. The final experiment used a similar toy dataset as the first experiment but now the first- and second-order moments were factored out and the model was trained to distinguish the different populations solely based on their skewness.

The toy dataset was created using the following steps. For each of the $n$ classes, a different low dimensional covariance matrix ($d \times d$) was created. Then, for each class, $m$ random $d$ dimensional points were generated from a standard normal distribution and rotated by calculating the dot product with their respective covariance matrix. By sampling all of the points from a standard normal distribution, all of their populations inherently have the same mean. The populations are then standardized, to make sure the model is learning the covariance and not the variance. Finally, $k$ samples of $q$ points are taken from each class, to simulate replicate populations as in the real data used in this study.

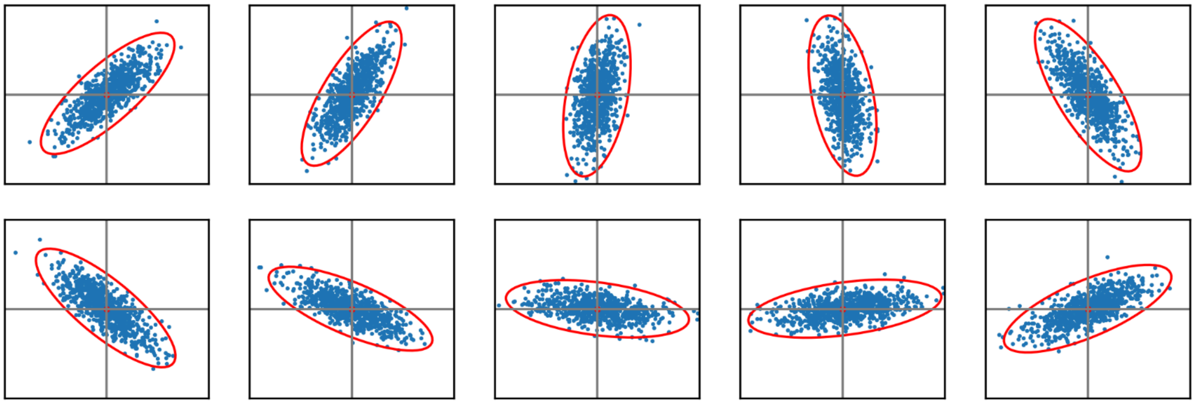

*Figure B1: Two-dimensional point set distributions for ten different population classes. The covariance distributions are indicated using red ellipsoids.*

For the first experiment, $n$, $d$, $m$, $q$, and $k$ are chosen to be 10, 2, 3200, 800, and 4 respectively, *Figure B1*. The training setup was similar to the setup used for the real single-cell feature data, which is described in the Experimental Setup. Six of the 10 classes were used to train the model and four were used as the test set. There was no need for a validation set, as no optimization for the model was done. The model architecture was generally the same as the one used for the real data, although the number of nodes per layer was drastically reduced. The input layer, first projection layer, and last projection layer consisted of 64, 32, and 2 nodes respectively. The optimizer and loss function were identical. The model was trained for 20 epochs with a batch size of 6.

The model was compared to the average-aggregated profile, which should give random results, by calculating the mAP for finding replicate populations in the test set. If the model is able to complete this task better than this baseline, that proves that this architecture is able to learn to infer covariance matrices from the input data. Although not irrefutable, it makes it more likely that the model for the single-cell feature data in this study is doing this as well.

After training the model in the described way, a mAP of 0.95 was achieved for finding replicate populations in the test set. This was higher than the baseline mAP of 0.31, which is similar to randomly choosing a sample in this test set. This means that the model is able to learn to infer the covariance matrices or some other higher-order statistic from the input data.

The second experiment aimed to verify these results by repeating the experiment on single-cell feature data from 11 Stain3 plates. Three plates were used as training plates and the eight other plates were used as a test set. The compounds were split in an 80% training and 20% validation set in the same way across all plates, just as described in the Experimental Setup. The per-well averages are subtracted from the cells, which means that the average profile should give random results again. The model is trained in the same way as in the main study, except for the number of epochs which was set to 40 instead of 100. The results are shown in *Table B1*.

The trained model achieved an average mAP of 0.30 and 0.23 across all plates on the training and validation compounds of the test set respectively. These were both higher than their respective baseline mAP of 0.03 and 0.04, which was random. This proves that the model is also able to learn to infer the covariance matrices or some other higher-order statistic from real high-dimensional input data.

The third experiment tested the degree to which the model can learn to infer second-order moments from the input data. In this experiment, the same toy dataset was used as in the first experiment, but now the populations are also gradually sphered. Sphering can be applied to a certain degree, using the regularization hyperparameter $r$. Here, a distinction is made between low ($r = 0.3$), medium ($r = 0.1$), and high ($r = 0.01$) sphering. Low sphering leaves much of the second-order moments intact, while high sphering removes nearly all of that information from the population. The mAP was calculated in the same way as in the first experiment. The model was trained in the same way as well, however, a smaller learning rate ($10^{-6}$ instead of $5 \times 10^{-4}$) and more epochs (1000 instead of 20) were required to train the model on this data. The different degrees of sphering are shown for one of the 10 classes in *Figure B2*.

*Table B1. Results of the second experiment. The mAP of the model and the baseline are shown for each training and test plate, and for the training and test compounds separately.*

| **plate** | **training compounds mAP model** | **training compounds mAP baseline** | **test compounds mAP model** | **test compounds mAP baseline** |
|---|---|---|---|---|
| *Training plates* | | | | |
| BR00115134 | 0.44 | 0.03 | 0.24 | 0.04 |
| BR00115125 | 0.37 | 0.03 | 0.25 | 0.04 |
| BR00115133highexp | 0.38 | 0.02 | 0.17 | 0.02 |
| *Test plates* | | | | |
| BR00115128highexp | 0.33 | 0.03 | 0.25 | 0.04 |
| BR00115125highexp | 0.29 | 0.03 | 0.22 | 0.02 |
| BR00115131 | 0.32 | 0.03 | 0.22 | 0.03 |
| BR00115133 | 0.32 | 0.03 | 0.16 | 0.04 |
| BR00115127 | 0.29 | 0.03 | 0.22 | 0.05 |

| | | | | |
|---|---|---|---|---|
| BR00115128 | 0.33 | 0.03 | 0.29 | 0.04 |
| BR00115129 | 0.3 | 0.03 | 0.26 | 0.04 |
| BR00115126 | 0.2 | 0.03 | 0.22 | 0.04 |

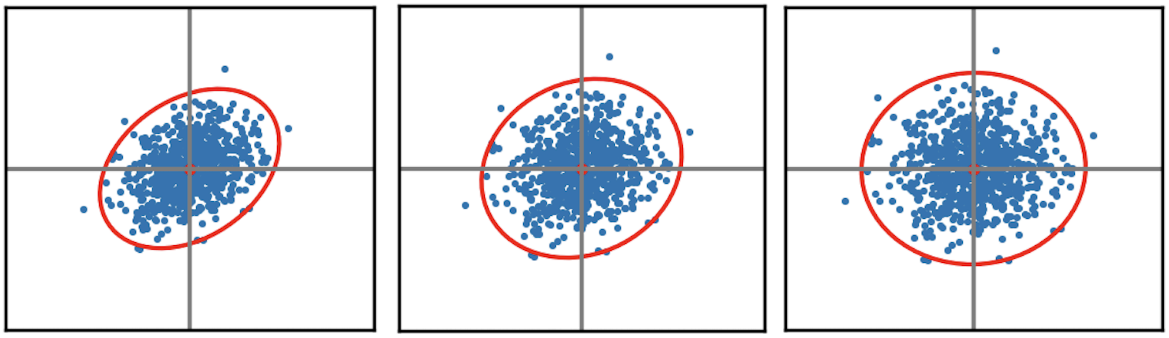

*Figure B2. Left: low sphering, middle: medium sphering, and right: heavy sphering on one of the 10 classes of the toy dataset.*

*Table B2. Results of the third experiment. The mAP of the model and baseline are shown for each degree of sphering, based on the sphering regularization parameter $r$.*

| **Degree of sphering** | **mAP model** | **mAP baseline** |
|---|---|---|
| low ($r = 0.3$) | 0.72 | 0.25 |
| medium ($r = 0.1$) | 0.61 | 0.25 |
| high ($r = 0.01$) | 0.29 | 0.25 |

The mAP of the model is about random after heavy sphering of the data, indicating that the model is no longer able to learn how to discern the different classes (Table B2). Even after varying a few hyperparameters like the model width, learning rate, and number of epochs the mAP did not improve. This is expected because the data can be almost fully explained by its second-order moments, which sphering factors out. It is not likely that this is the case for real single-cell feature data, where more interactions are expected.

After medium or low sphering, the model is still able to beat the baseline, although it requires many more training steps and a smaller learning rate than in the first experiment. In fact, the degree to which second-order moments are present correlates with the mAP achieved by the model.

The final experiment tested the degree to which the model can learn to infer third-order moments from the input data. A toy dataset was created in a similar way to the first experiment (Figure B3). For each of the $n$ classes, a skewness value was sampled. Then, for each class, $m$ random $1$ dimensional points were generated from a standard normal distribution and skewed according to their respective skewness value. The populations are then standardized. Finally, $k$ samples of $q$ points are taken from each class, to simulate replicate populations as in the real data used in this study. The same values were used for $n$, $d$, $m$, $q$, and $k$ as in the first experiment, except for $d$ which is now chosen to be 1.

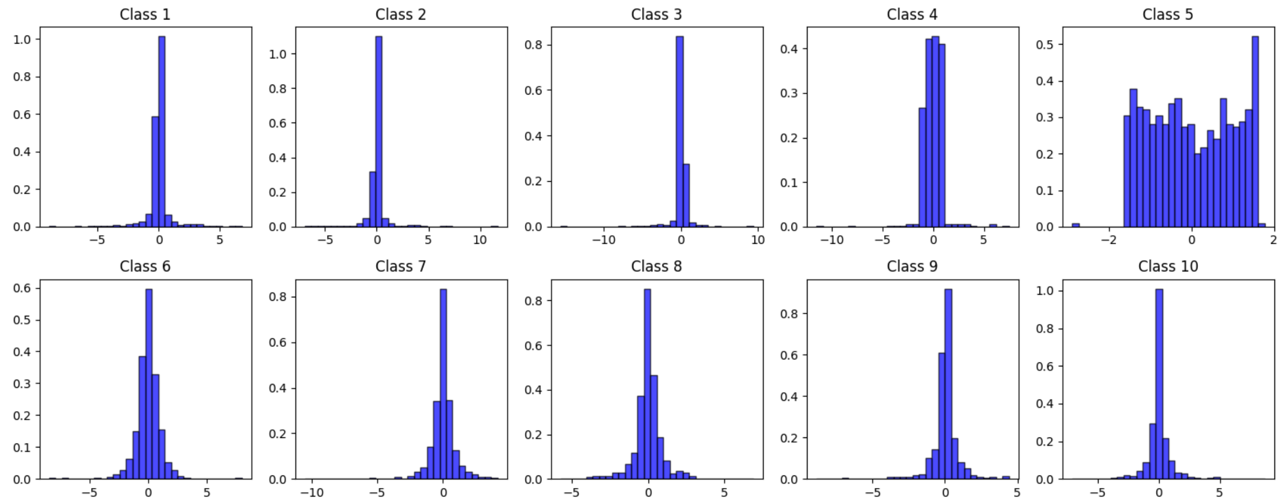

*Figure B3: One-dimensional point set distributions for ten different population classes. The mean and standard deviation of all classes are 0 and 1, respectively.*

The same stratification as the first experiment was used: six of the 10 classes were used to train the model and four were used as the test set. After training the model (with the same setup as before), a mAP of 0.34 was achieved for finding replicate populations in the test set. This was higher than the baseline mAP of 0.22, which is similar to randomly choosing a sample in this test set. The improvement is not as pronounced as we saw for learning to infer the covariance from the input data. This could be due to the task being more difficult, the limitation of having only one feature dimension, or a combination thereof. Nevertheless, we can reasonably assume that the model is able to learn to infer third-order moments from the input data in addition to second-order moments.

These experiments show that the model is indeed learning to infer second- and third-order moments from the input data. The different experiments also quantify the complexity of the task as the number of epochs or learning rate had to be adjusted to properly train the model. It was easier for the model to learn second-order moments from zero-meaned and standardized two dimensional feature data than high dimensional ($d = 1324$) feature data. These findings were reaffirmed by the final experiment where we observed a relatively small improvement in mAP of the model compared to random chance. Additionally, as the presence of the second-order moments was gradually decreased in the input data the model's ability to learn gradually decreased as well.

### C. Training, validation, and test set stratification

*Table C1: The training, validation, and test set stratification for Stain2, Stain3, Stain4, and Stain5. Five training, four validation, and three test plates are used for Stain2, Stain3, and Stain4. Stain5 contains six test set plates only. The plate metadata can be found here [https://github.com/carpenter-singh-lab/2023_Cimini_NatureProtocols/blob/7308631ee936669c58397d6dd0d3dd6d9745eea9/JUMPExperimentMasterTable.csv](https://github.com/carpenter-singh-lab/2023_Cimini_NatureProtocols/blob/7308631ee936669c58397d6dd0d3dd6d9745eea9/JUMPExperimentMasterTable.csv).*

| Stain2 | Stain3 | Stain4 | Stain5 |
|---|---|---|---|
| *Training plates* | | | *Test plates* |
| BR00113818 | BR00115128 | BR00116627 | BR00120532 |
| BR00113820 | BR00115125highexp | BR00116631 | BR00120270 |
| BR00112202 | BR00115133highexp | BR00116625 | BR00120536 |
| BR00112197binned | BR00115131 | BR00116630highexp | BR00120530 |
| BR00112198 | BR00115134 | 200922_015124-Vhighexp | BR00120526 |
| *Validation plates* | | | BR00120274 |
| BR00112197standard | BR00115129 | BR00116628highexp | |
| BR00112197repeat | BR00115133 | BR00116629highexp | |
| BR00112204 | BR00115128highexp | BR00116627highexp | |
| BR00112201 | BR00115127 | BR00116629 | |
| *Test plates* | | | |
| BR00112199 | BR00115134bin1 | 200922_044247-Vbin1 | |
| BR00113819 | BR00115134multiplane | 200922_015124-V | |
| BR00113821 | BR00115126highexp | BR00116633bin1 | |

### D. CytoSummaryNet hyperparameter overview

We used the Weights & Biases (WandB) sweep suite in combination with the BOHB (Bayesian Optimization and HyperBand) algorithm for hyperparameter sweeps. The BOHB algorithm [47] combines Bayesian optimization with bandit-based strategies to efficiently find optimal hyperparameters.

Additionally Table D1 provides an overview of all tunable hyperparameters and their chosen values based on a BOHB hyperparameter optimization.

*Table D1: Overview of all tunable hyperparameters and their chosen values based on a random search hyperparameter optimization.*

| Hyperparameter | cpg0001 value | cpg0004 value |
|---|---|---|
| AdamW learning rate | $5 \times 10^{-4}$ | $5 \times 10^{-4}$ |
| AdamW weight decay | $10^{-2}$ | $10^{-2}$ |
| epochs | 100 | 100 |
| number of different compounds per batch | *18* | *32* |

| | | |
|---|---|---|
| number of samples per compound | *4* | *8* |
| batch size | $18 \times 4 = 72$ | $32 \times 8 = 256$ |
| Gaussian mean (for sampling number of cells) | $1500$ | $1500$ |
| Gaussian standard deviation (for sampling number of cells) | $800$ | $800$ |
| latent dimension after first layer | $2048$ | $2048$ |
| latent dimension after first projection layer | $256$ | $256$ |
| number of projection layers | 2 | 2 |
| output dimension of model (loss/aggregated profile space) | $2048$ | $2048$ |
| SupCon loss temperature | $0.1$ | $0.1$ |

### E. Relevance correlation with CellProfiler features

*Table E1: Top 20 CellProfiler features based on their negative Pearson correlation coefficient with the SA and CPA combined relevance score. The scores were calculated for a single test plate of Stain3 (200922_015124-V).*

| Feature category | Feature name | Correlation |
|---|---|---|
| Cells | Cells_Intensity_MeanIntensityEdge_DNA | -0.74 |
| Cytoplasm | Cytoplasm_Intensity_MeanIntensityEdge_DNA | -0.72 |
| Cytoplasm | Cytoplasm_Intensity_UpperQuartileIntensity_DNA | -0.71 |
| Cytoplasm | Cytoplasm_Intensity_MeanIntensity_DNA | -0.69 |
| Cytoplasm | Cytoplasm_Correlation_K_Brightfield_DNA | -0.67 |
| Cells | Cells_Intensity_MedianIntensity_DNA | -0.64 |
| Cells | Cells_Intensity_MeanIntensity_DNA | -0.63 |
| Cells | Cells_Intensity_MeanIntensityEdge_ER | -0.61 |
| Cytoplasm | Cytoplasm_Correlation_K_Mito_DNA | -0.61 |
| Cytoplasm | Cytoplasm_Intensity_StdIntensity_DNA | -0.61 |
| Cytoplasm | Cytoplasm_Intensity_MedianIntensity_DNA | -0.61 |
| Cytoplasm | Cytoplasm_Intensity_MADIntensity_DNA | -0.61 |
| Cells | Cells_Intensity_MeanIntensityEdge_RNA | -0.6 |
| Cells | Cells_Intensity_MeanIntensityEdge_AGP | -0.6 |
| Cytoplasm | Cytoplasm_Intensity_MeanIntensityEdge_RNA | -0.6 |
| Cytoplasm | Cytoplasm_Intensity_MeanIntensityEdge_ER | -0.59 |
| Cytoplasm | Cytoplasm_RadialDistribution_RadialCV_DNA_3of4 | -0.58 |
| Cells | Cells_Intensity_LowerQuartileIntensity_DNA | -0.58 |
| Cells | Cells_Intensity_MaxIntensityEdge_RNA | -0.58 |
| Cells | Cells_Intensity_MaxIntensityEdge_ER | -0.57 |

*Table E2: Top 20 CellProfiler features based on their negative Pearson correlation coefficient with the SA and CPA combined relevance score. The scores were calculated for a single test plate of Stain3 (200922_015124-V).*

| Feature category | Feature name | Correlation |
|---|---|---|
| Cytoplasm | Cytoplasm_Correlation_K_DNA_Brightfield | 0.72 |
| Cells | Cells_AreaShape_MeanRadius | 0.71 |
| Cells | Cells_AreaShape_MaximumRadius | 0.7 |
| Cells | Cells_AreaShape_MedianRadius | 0.7 |
| Cells | Cells_AreaShape_Area | 0.68 |
| Cells | Cells_AreaShape_MinorAxisLength | 0.68 |

| | | |
|---|---|---|
| Cells | Cells_AreaShape_MinFeretDiameter | 0.68 |
| Cytoplasm | Cytoplasm_AreaShape_MinFeretDiameter | 0.68 |
| Cytoplasm | Cytoplasm_Intensity_IntegratedIntensityEdge_Brightfield | 0.68 |
| Cells | Cells_Intensity_IntegratedIntensity_Brightfield | 0.67 |
| Cytoplasm | Cytoplasm_AreaShape_Perimeter | 0.67 |
| Cytoplasm | Cytoplasm_AreaShape_MinorAxisLength | 0.66 |
| Cytoplasm | Cytoplasm_AreaShape_Area | 0.65 |
| Cytoplasm | Cytoplasm_Intensity_IntegratedIntensity_Brightfield | 0.65 |
| Cells | Cells_AreaShape_Perimeter | 0.64 |
| Nuclei | Nuclei_AreaShape_MedianRadius | 0.64 |
| Cells | Cells_Intensity_IntegratedIntensityEdge_Brightfield | 0.64 |
| Nuclei | Nuclei_AreaShape_MeanRadius | 0.64 |
| Cytoplasm | Cytoplasm_AreaShape_MedianRadius | 0.64 |
| Cytoplasm | Cytoplasm_AreaShape_MeanRadius | 0.64 |

*Table E3: Top 20 CellProfiler features based on their negative Pearson correlation coefficient with the SA relevance score. The scores were calculated for a single test plate of Stain3 (200922_015124-V).*

| Feature category | Feature name | Correlation |
|---|---|---|
| Cells | Cells_Intensity_MeanIntensityEdge_DNA | -0.69 |
| Cytoplasm | Cytoplasm_Intensity_MeanIntensityEdge_DNA | -0.67 |
| Cytoplasm | Cytoplasm_Intensity_UpperQuartileIntensity_DNA | -0.67 |
| Cytoplasm | Cytoplasm_Correlation_K_Brightfield_DNA | -0.66 |
| Cytoplasm | Cytoplasm_Intensity_MeanIntensity_DNA | -0.66 |
| Cells | Cells_Intensity_MeanIntensity_DNA | -0.61 |
| Cells | Cells_Intensity_MedianIntensity_DNA | -0.60 |
| Cells | Cells_Intensity_MaxIntensityEdge_RNA | -0.58 |
| Cytoplasm | Cytoplasm_Intensity_MeanIntensityEdge_RNA | -0.58 |
| Cytoplasm | Cytoplasm_Intensity_MADIntensity_DNA | -0.58 |
| Cytoplasm | Cytoplasm_Intensity_MedianIntensity_DNA | -0.58 |
| Cells | Cells_Intensity_MaxIntensityEdge_ER | -0.58 |
| Cells | Cells_Intensity_MeanIntensityEdge_ER | -0.58 |
| Cytoplasm | Cytoplasm_Intensity_MeanIntensityEdge_ER | -0.58 |
| Cytoplasm | Cytoplasm_Intensity_StdIntensity_DNA | -0.57 |
| Cells | Cells_Intensity_MeanIntensityEdge_AGP | -0.57 |
| Cells | Cells_Intensity_MeanIntensityEdge_RNA | -0.57 |
| Cytoplasm | Cytoplasm_Intensity_MeanIntensityEdge_AGP | -0.56 |
| Cytoplasm | Cytoplasm_RadialDistribution_RadialCV_DNA_3of4 | -0.56 |
| Nuclei | Nuclei_RadialDistribution_RadialCV_DNA_4of4 | -0.55 |

*Table E4: Top 20 CellProfiler features based on their positive Pearson correlation coefficient with the SA relevance score. The scores were calculated for a single test plate of Stain3 (200922_015124-V).*

| Feature category | Feature name | Correlation |
|---|---|---|
| Cytoplasm | Cytoplasm_Correlation_K_DNA_Brightfield | 0.68 |
| Cells | Cells_AreaShape_MeanRadius | 0.63 |
| Cells | Cells_AreaShape_MaximumRadius | 0.62 |
| Cells | Cells_AreaShape_MedianRadius | 0.62 |
| Cells | Cells_AreaShape_MinorAxisLength | 0.60 |
| Cytoplasm | Cytoplasm_AreaShape_MinFeretDiameter | 0.60 |
| Cells | Cells_AreaShape_MinFeretDiameter | 0.60 |
| Cytoplasm | Cytoplasm_Intensity_IntegratedIntensityEdge_Brightfield | 0.59 |
| Nuclei | Nuclei_AreaShape_MedianRadius | 0.58 |
| Cells | Cells_AreaShape_Area | 0.58 |
| Nuclei | Nuclei_AreaShape_MeanRadius | 0.58 |
| Cells | Cells_Intensity_IntegratedIntensity_Brightfield | 0.58 |
| Cytoplasm | Cytoplasm_AreaShape_Perimeter | 0.58 |

| | | |
|---|---|---|
| Cytoplasm | Cytoplasm_AreaShape_MinorAxisLength | 0.57 |
| Cytoplasm | Cytoplasm_AreaShape_MeanRadius | 0.57 |
| Cells | Cells_Neighbors_FirstClosestDistance_Adjacent | 0.57 |
| Cytoplasm | Cytoplasm_AreaShape_MedianRadius | 0.57 |
| Cytoplasm | Cytoplasm_AreaShape_MaximumRadius | 0.56 |
| Cells | Cells_Intensity_IntegratedIntensityEdge_Brightfield | 0.56 |
| Cytoplasm | Cytoplasm_AreaShape_Area | 0.56 |

*Table E5: Top 20 CellProfiler features based on their negative Pearson correlation coefficient with the CPA relevance score. The scores were calculated for a single test plate of Stain3 (200922_015124-V).*

| **Feature category** | **Feature name** | **Correlation** |
|---|---|---|
| Cytoplasm | Cytoplasm_Correlation_K_Mito_DNA | -0.51 |
| Cells | Cells_Intensity_MeanIntensityEdge_DNA | -0.49 |
| Cells | Cells_Correlation_Overlap_Mito_RNA | -0.48 |
| Cytoplasm | Cytoplasm_Intensity_MeanIntensityEdge_DNA | -0.48 |
| Cells | Cells_RadialDistribution_MeanFrac_Mito_4of4 | -0.47 |
| Cytoplasm | Cytoplasm_Correlation_Overlap_Mito_RNA | -0.47 |
| Cytoplasm | Cytoplasm_Correlation_K_Mito_AGP | -0.46 |
| Cytoplasm | Cytoplasm_Correlation_Manders_Brightfield_Mito | -0.46 |
| Cells | Cells_Correlation_Manders_Brightfield_Mito | -0.46 |
| Cells | Cells_Correlation_Overlap_AGP_Brightfield | -0.46 |
| Cytoplasm | Cytoplasm_Correlation_Manders_RNA_Mito | -0.45 |
| Cytoplasm | Cytoplasm_Correlation_Manders_ER_Mito | -0.45 |
| Cytoplasm | Cytoplasm_Correlation_Manders_AGP_Mito | -0.45 |
| Cells | Cells_Correlation_Manders_ER_Mito | -0.45 |
| Cells | Cells_Correlation_Manders_RNA_Mito | -0.45 |
| Cytoplasm | Cytoplasm_Correlation_K_Mito_RNA | -0.45 |
| Cells | Cells_Correlation_Manders_AGP_Mito | -0.45 |
| Cytoplasm | Cytoplasm_Correlation_Manders_DNA_Mito | -0.45 |
| Cells | Cells_Correlation_K_Mito_AGP | -0.45 |
| Cytoplasm | Cytoplasm_Intensity_UpperQuartileIntensity_DNA | -0.45 |

*Table E6: Top 20 CellProfiler features based on their positive Pearson correlation coefficient with the CPA relevance score. The scores were calculated for a single test plate of Stain3 (200922_015124-V).*

| **Feature category** | **Feature name** | **Correlation** |
|---|---|---|
| Cells | Cells_Intensity_IntegratedIntensity_Mito | 0.67 |
| Cytoplasm | Cytoplasm_Intensity_IntegratedIntensity_Mito | 0.65 |
| Cytoplasm | Cytoplasm_Intensity_IntegratedIntensityEdge_Mito | 0.60 |
| Cells | Cells_AreaShape_Area | 0.59 |
| Cells | Cells_Intensity_IntegratedIntensity_Brightfield | 0.58 |
| Cytoplasm | Cytoplasm_AreaShape_Area | 0.57 |
| Cells | Cells_Intensity_IntegratedIntensity_AGP | 0.57 |
| Cytoplasm | Cytoplasm_Intensity_IntegratedIntensity_Brightfield | 0.56 |
| Nuclei | Nuclei_Intensity_IntegratedIntensityEdge_Mito | 0.56 |
| Cytoplasm | Cytoplasm_AreaShape_Perimeter | 0.56 |
| Cytoplasm | Cytoplasm_Intensity_IntegratedIntensity_DNA | 0.56 |
| Cells | Cells_AreaShape_MaximumRadius | 0.55 |
| Cells | Cells_Intensity_StdIntensity_Mito | 0.55 |
| Cells | Cells_AreaShape_MeanRadius | 0.55 |
| Cytoplasm | Cytoplasm_Intensity_IntegratedIntensity_AGP | 0.55 |
| Cytoplasm | Cytoplasm_Intensity_IntegratedIntensityEdge_Brightfield | 0.55 |
| Cytoplasm | Cytoplasm_Correlation_K_DNA_Mito | 0.55 |
| Cells | Cells_AreaShape_MedianRadius | 0.55 |
| Cells | Cells_AreaShape_Perimeter | 0.55 |
| Cells | Cells_Intensity_IntegratedIntensity_DNA | 0.54 |

#### F. Measuring plate similarity

Plate similarity is measured using a hierarchical clustering analysis. First, the average-aggregated profile is taken per well for each available plate. This results in 384 well profiles of 1324 features per plate. PCA analysis is then performed on this matrix and the loadings of the first principal component (PC1) are taken, resulting in a 1x1324 vector. This vector is then normalized to a unit vector. These vectors are calculated for all available plates and the Pearson correlation is calculated between them. Comparing the PC1 loadings of two multivariate distributions is a very rough approximation for comparing their covariance matrices [48], and thus the Pearson correlation can quantify their similarity. Finally, the Pearson correlation is used to create a hierarchical clustering map, Figure F1. The clustermap shows that Stain5 and Stain2, Stain3, and Stain4 can be separated as the two main clusters, signifying the strong experiment effects between these two groups. We chose the plates that are most distinct and dissimilar to the rest of the plates within each cluster for Stain2, Stain3, and Stain4 to be test plates.

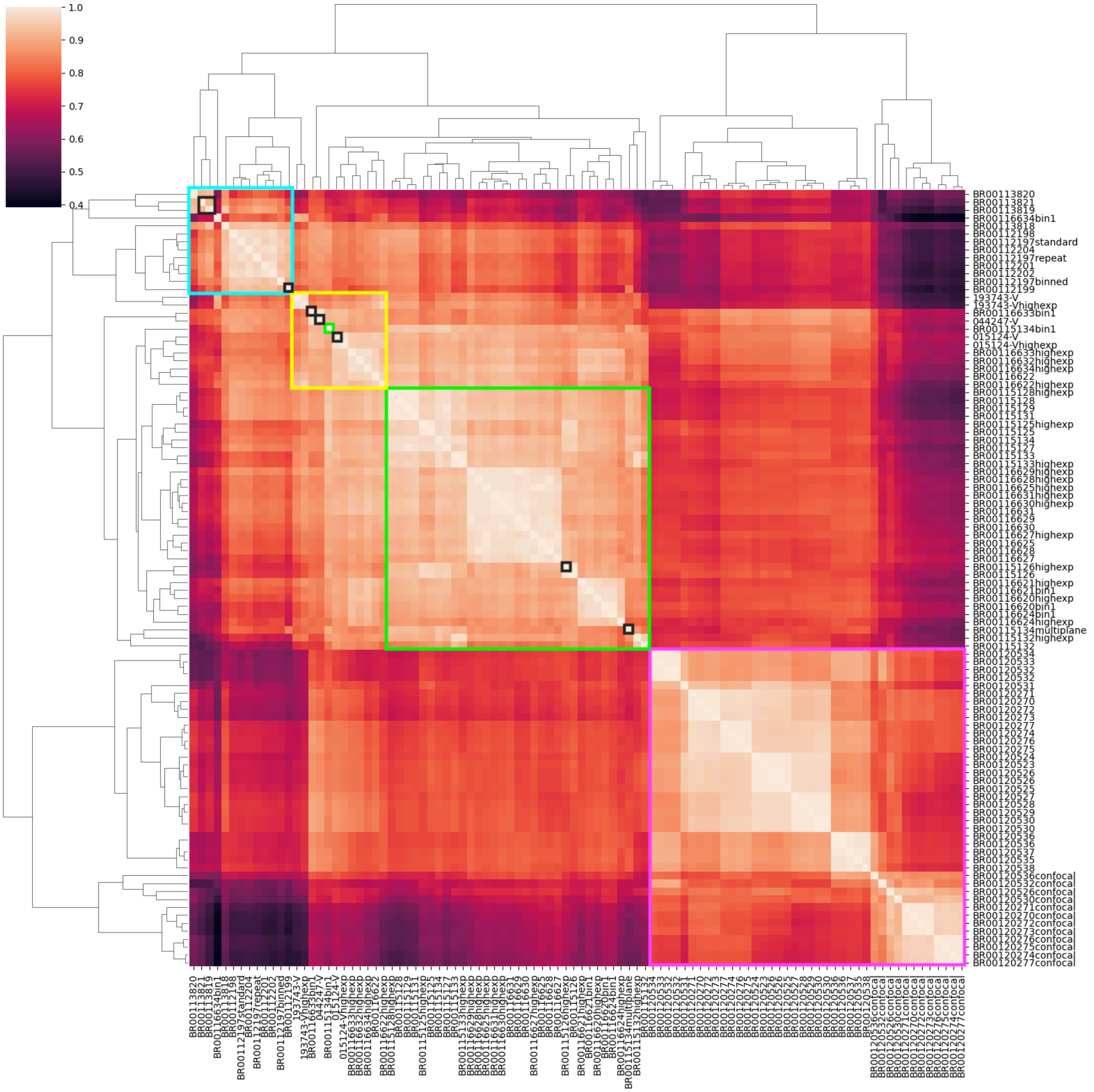


Figure F1: Hierarchical clustermap of the Pearson correlations between the PC1 loadings of the mean aggregated profile features per plate. The plate clusters of Stain2 (cyan), Stain4 (yellow), Stain3 (green), and Stain5 (pink) are annotated with boxes. The test plates within each StainX subset are annotated with black boxes, one test plate from Stain3, annotated in green, which is an outlier and falls within the Stain4 cluster. All plates of Stain5 are also test plates, but not annotated to reduce visual clutter.

#### G. Additional cell image FOVs

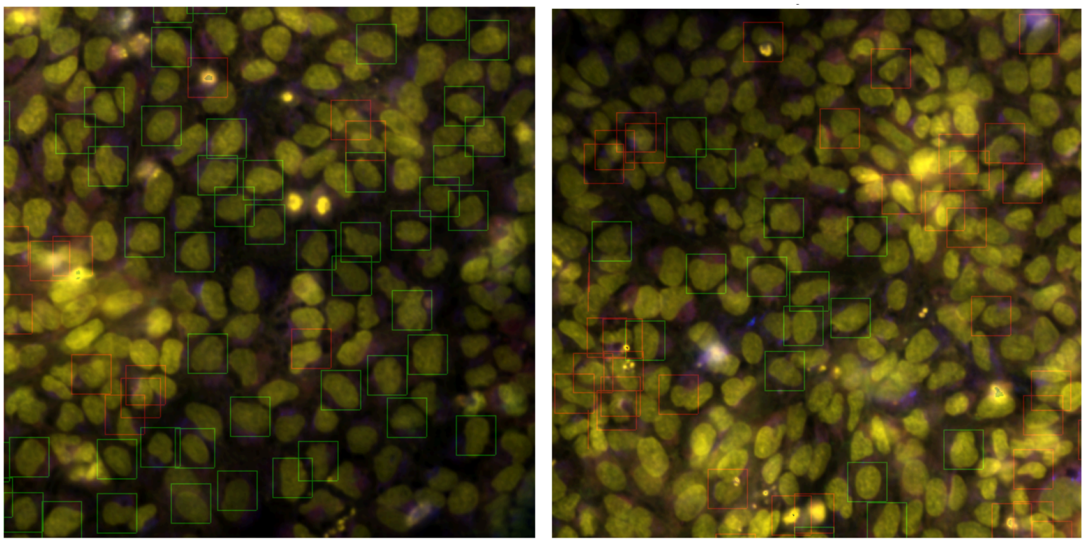

Figure G1: 5-channel combined microscope image of the second and third FOVs for plate BR00112197binned. The most relevant cells are annotated with green boxes and the least relevant cells are annotated with red boxes.

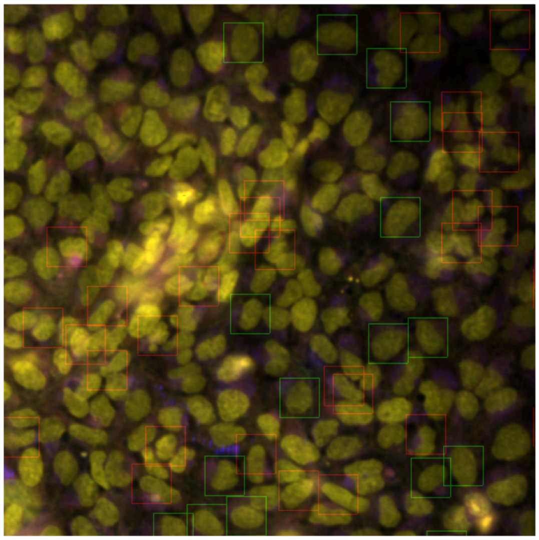

Figure G2: 5-channel combined microscope image of the fourth FOV for plate BR00112197binned. The most relevant cells are annotated with green boxes and the least relevant cells are annotated with red boxes.

### H. Layout of training and validation compounds

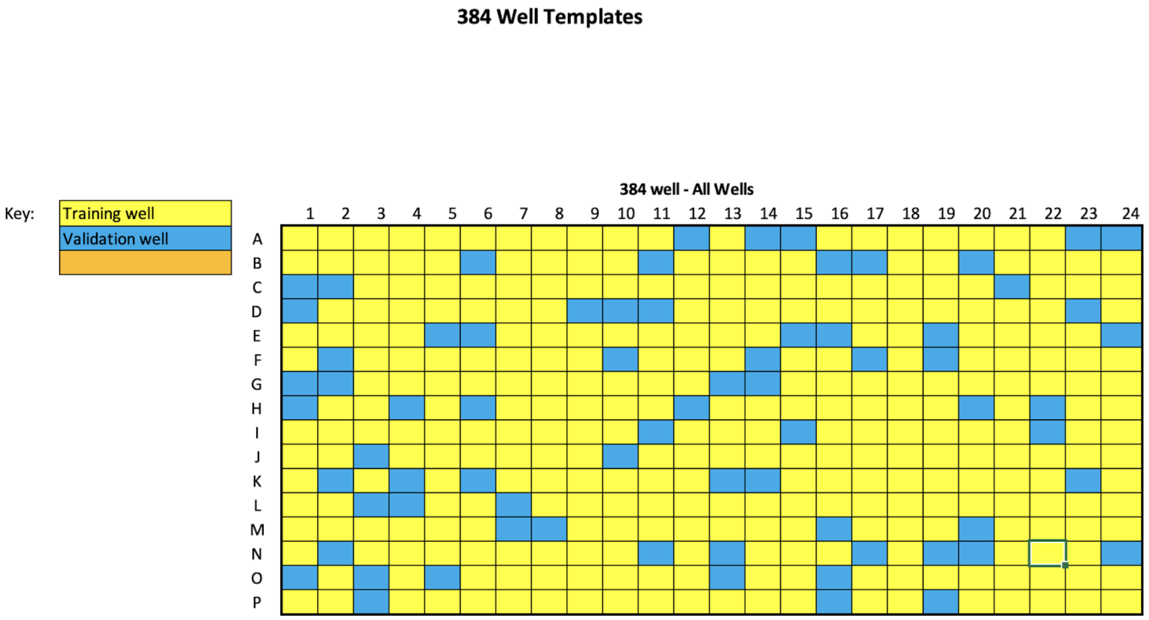


*Figure H1: Training and validation compound split for all plates of cpg0001. The compounds were selected at random.*

### I. Relevant features for CytoSummaryNet improvement

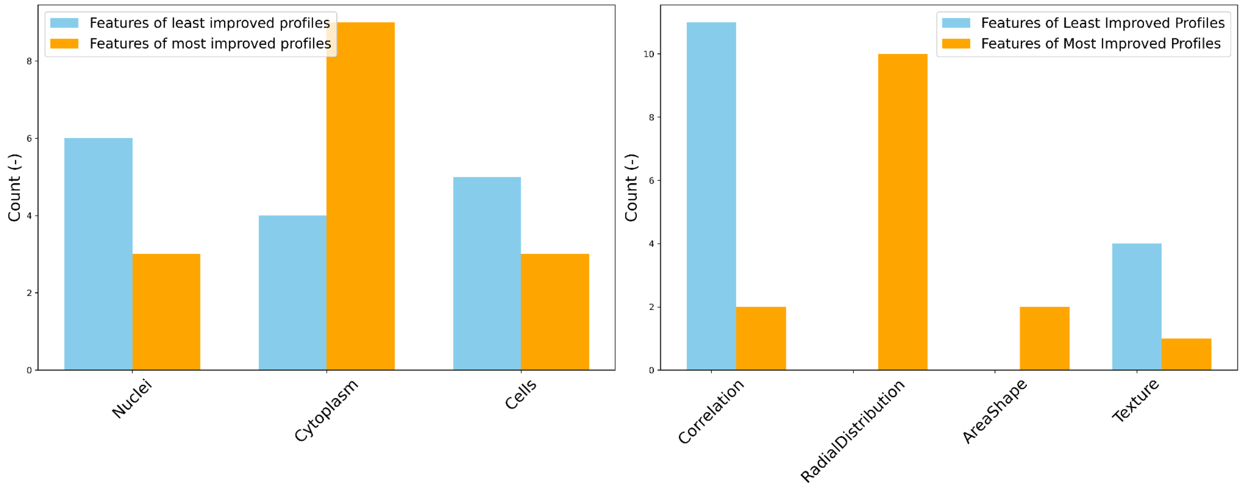


*Figure I1: Distribution of feature types that most strongly differentiate average profiles from negative controls across mechanisms of action most (orange) and least (lightblue) improved by CytoSummaryNet as compared to average profiling. (Left) Distribution of feature compartment (Nuclei, Cells, or Cytoplasm); (right) distribution of feature type.*

*Table I1: List of features that most strongly differentiate average profiles from negative controls, for a subset of mechanisms of action (MoA).*

| MoA | Top 5 distinct features from negative controls | mAP average/CSN |
|---|---|---|
| JNK inhibitor | Cytoplasm_**RadialDistribution**_RadialCV_**Mito**_3of4<br>Cells_**RadialDistribution**_RadialCV_**Mito**_4of4<br>Cytoplasm_**RadialDistribution**_RadialCV_**Mito**_2of4<br>Cells_**Correlation**_Overlap_**Mito**_Brightfield<br>Cytoplasm_**Correlation**_Overlap_**Mito**_Brightfield | 0.48 / 0.96 |
| mTOR inhibitor | Nuclei_**AreaShape**_Zernike_2_0<br>Nuclei_**Texture**_InverseDifferenceMoment_Brightfield_5_00<br>Cytoplasm_**RadialDistribution**_FracAtD_**Mito**_2of4<br>Cytoplasm_**RadialDistribution**_FracAtD_**Mito**_1of4<br>Nuclei_**AreaShape**_Zernike_5_3 | 0.40 / 0.79 |
| inosine monophosphate dehydrogenase inhibitor | Cytoplasm_**RadialDistribution**_MeanFrac_**RNA**_1of4<br>Cytoplasm_**RadialDistribution**_MeanFrac_**RNA**_4of4<br>Cytoplasm_**RadialDistribution**_FracAtD_**RNA**_2of4<br>Cells_**RadialDistribution**_RadialCV_**RNA**_2of4<br>Cytoplasm_**RadialDistribution**_FracAtD_**RNA**_1of4 | 0.03 / 0.84 |
| EGFR inhibitor | Cytoplasm_**Correlation**_Costes_RNA_**Brightfield**<br>Nuclei_**Correlation**_Manders_**Brightfield**_AGP<br>Nuclei_**Correlation**_Manders_DNA_AGP<br>Cells_**Correlation**_Costes_AGP_**Brightfield**<br>Nuclei_**Correlation**_Costes_**Brightfield**_AGP | 0.33 / 0.31 |
| BCL inhibitor | Cytoplasm_**Correlation**_Costes_ER_**Brightfield**<br>Cells_**Correlation**_Costes_**Brightfield**_ER<br>Nuclei_**Correlation**_Costes_DNA_**Brightfield**<br>Cells_**Correlation**_Costes_**Brightfield**_Mito<br>Nuclei_**Correlation**_Costes_**Brightfield**_DNA | 0.12 / 0.11 |
| hypoxia inducible factor inhibitor | Cells_**Texture**_InfoMeas2_ER_20_01<br>Cytoplasm_**Texture**_InfoMeas2_ER_20_00<br>Cytoplasm_**Texture**_InfoMeas2_ER_20_02<br>Cells_**Texture**_InfoMeas2_ER_20_03<br>Nuclei_**Correlation**_Manders_**Brightfield**_AGP | 0.39 / 0.29 |

### J. Computation time and storage requirements comparison

*Table J1: Comparison of computation time and storage requirements between CytoSummaryNet and average profiling. The table lists the average computation times and storage sizes for various stages of both methods. CytoSummaryNet profiling adds computation time and storage requirements during the aggregation process compared to average profiling. CytoSummaryNet training requires an intermediate form of single-cell data storage for quick access and takes one to two days, depending on the type of CPU or GPU used; we used an Intel(R) Xeon(R) Platinum 8375C CPU @ 2.90GHz and a 2.4 GHz 8-Core Intel Core i9, respectively. The intermediate data storage size for CytoSummaryNet is a fraction (0.44) of the original raw SQLite file, due in part to the conversion of double-precision numbers to single-precision numbers. This conversion also contributes to the reduction in the storage space required for the final aggregated profiles when using CytoSummaryNet aggregation compared to average profiling.*

| method | preprocessing time per plate (seconds) | intermediate data storage size per plate (GB per raw SQLite GB) | model training time (hours) | profile aggregation computation time per plate (seconds) | csv file size of aggregated profiles per plate (MB) |
|---|---|---|---|---|---|
| CytoSummaryNet profiling | ~50 | 0.44 | ~26 to ~51* | 20.1 | ~8 |
| average profiling | ~40 | N/A | N/A | 0.4 | ~30 |

#### K. Number of predictable mechanisms of action across different mAP thresholds

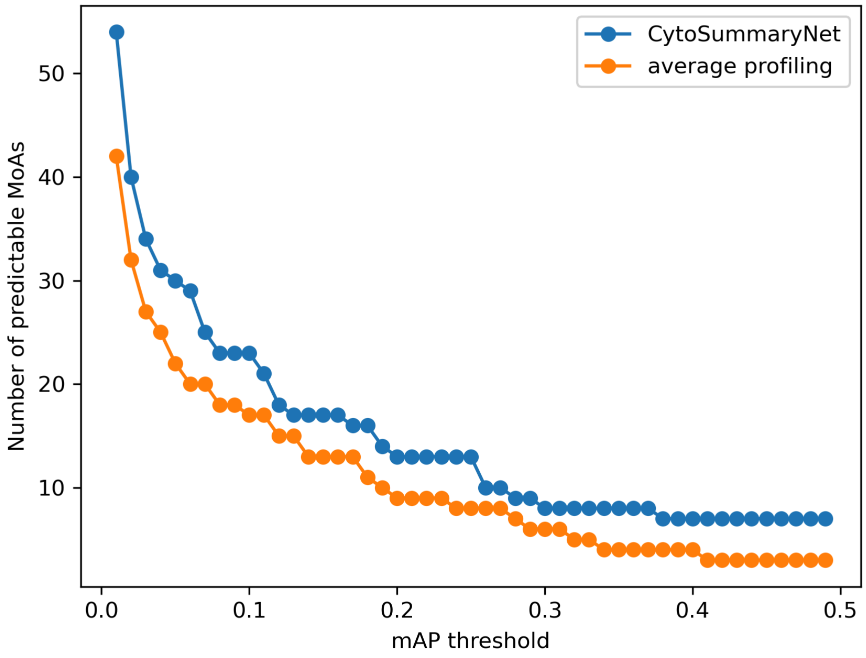


*Figure K1: Number of predictable mechanisms of action (MoAs) for cpg0004 profiles created by CytoSummaryNet (blue) and average profiling (orange) as a function of an increasing mAP threshold. This is an alternative method of representing the data in Figure 6.*

### L. Investigating CytoSummaryNet's cell prioritization

To validate our interpretation that CytoSummaryNet prioritizes large, uncrowded cells, we conducted an additional analysis using five different plates from LINCS at the 10 µM dose point, resulting in approximately 50 wells to compute the replicate retrieval mAP.

We progressively filtered out cells based on a quantile threshold for Cells_AreaShape features (MeanRadius, MaximumRadius, MedianRadius, and Area), which were identified as important in our interpretability analysis, and then computed average profiles using the remaining cells before determining the replicate retrieval mAP. In the exclusion experiment (Figure L1; left), we gradually left out cells as the threshold increased from 0.0 (all cells included) to 1.0 (only the largest cells included). In the inclusion experiment (Figure L1; right), we progressively included larger cells from left to right, with the leftmost point representing no cells and the rightmost point representing all cells.

The results show that using only the largest cells does not significantly increase the performance compared to using all cells (rightmost points in both figures). Instead, the key finding is that including large cells is more important than only including small cells. The mAP saturates after a threshold of around 0.4 in both experiments, indicating that larger cells define the profile the most, and once enough cells are included to outweigh the smaller cell features, the profile does not change significantly by including even larger cells.

These findings support our interpretation that CytoSummaryNet prioritizes large, uncrowded cells. While this approach could potentially be used as a general outlier removal strategy for cell profiling, further investigation is needed to assess its robustness and generalizability across different datasets and experimental conditions.

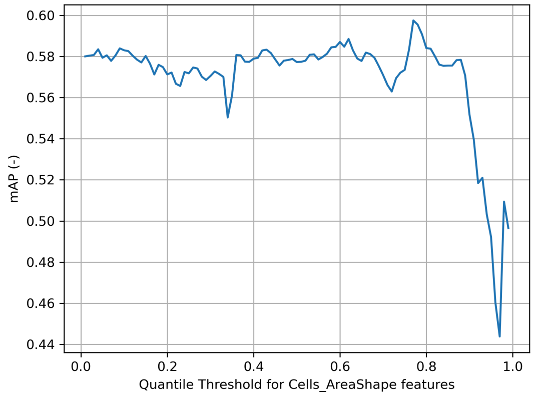


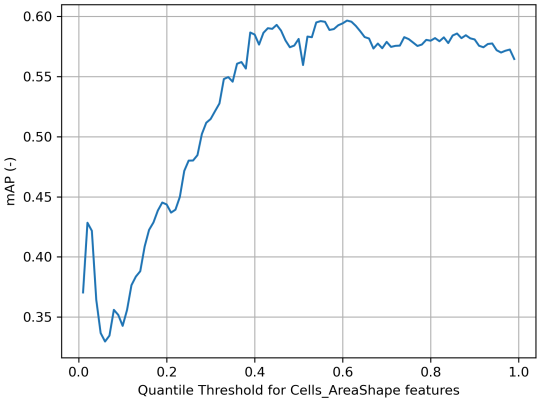


*Figure L1. The impact of progressively (left) excluding, and (right) including cells based on their Cells_AreaShape features on the replicate retrieval mAP. The x-axis represents the quantile threshold, with 0.0 including all cells and 1.0 including only the largest cells. The y-axis represents the mAP score.*
